# Repetitive Learning Control for Body Caudal Undulation with Soft Sensory Feedback

**DOI:** 10.1101/2024.01.11.575004

**Authors:** Fabian Schwab, Mohamed El Arayshi, Seyedreza Rezaei, Hadrien Sprumont, Federico Allione, Claudio Mucignat, Ivan Lunati, Cristiano Maria Verrelli, Ardian Jusufi

**Affiliations:** Soft Kinetic Group, Engineering Sciences Department, Swiss Federal Laboratories for Materials Science and Technology, Dübendorf, Switzerland; Locomotion in Biorobotic and Somatic Systems Group, Max Planck Institute for Intelligent Systems, Germany; Department of Electronic Engineering, University of Rome Tor Vergata, Italy; Laboratory for Computational Engineering, Swiss Federal Laboratories for Materials Science and Technology, Dübendorf, Switzerland; Institute for NeuroInformatics, ETH and University of Zürich, Switzerland

**Author notes:** Correspondence: Ardian Jusufi.

**Keywords:** Bio-inspired, Soft robotics, Repetitive Learning, Robotic fish, Locomotion, Swimming, Soft sensor, Experimental validation

## Abstract

Soft bio-inspired robotics is a growing field of research that seeks to close the gap with animal robustness and adaptability where conventional robots fall short. The embedding of sensors with the capability to discriminate between different body deformation modes is a key technological challenge in soft robotics to enhance robot control – a difficult task for such kinds of systems with high degrees of freedom. The recently conceived Linear Repetitive Learning Estimation Scheme (LRLES) – to be included in the traditional Proportional Integral Derivative (PID) control – is proposed here as a way to compensate for uncertain dynamics on a soft swimming robot, which is actuated with soft pneumatic actuators and equipped with soft sensors providing proprioceptive information pertaining to lateral body caudal bending akin to a goniometer. The proposed controller is derived in detail and experimentally validated, with the experiment consisting of tracking a desired trajectory for bending angle while continuously oscillating with a constant frequency. The results are compared vis a vis those achieved with the traditional PID controller, finding that the PID endowed with the LRLES outperforms the PID controller (though the latter has been separately tuned) and experimentally validating the novel controller’s effectiveness, accuracy, and matching speed.

## 1 INTRODUCTION

The last two decades accelerated a paradigm shift in experimental robotics, driven in part by computational advances as well as by the comparative analysis of biological systems. This shift has catalyzed the development of sophisticated robots that are borrow from general principles of biomechanics and neurocontrol of living organisms, as delineated in the foundational works of (Dickinson et al., 2000; Ijspeert, 2020). Despite remarkable progress in crafting robots that mirror the form and function of biological entities, a significant gap remains in replicating the adaptability and resilience inherent in natural locomotion. This disparity is particularly evident in areas such as sensory perception and adaptive response mechanisms. The unparalleled dexterity of animals, exemplified by geckos maneuvering on water surfaces (Nirody et al., 2018), crocodiles in their prey-capture patterns (Fish et al., 2007), and the enduring resilience of migratory fish (Crossin et al., 2004), underscores the profound complexity and efficiency of biological systems. This gap primarily arises from animals’ flexible anatomies combined with integrated sensing, which grants them an ability termed ‘morphological intelligence’ (Woodward and Sitti, 2018; Martinez-Hernandez et al., 2019).

The objectives of bioinspired robotics encompass a triad of goals: to decode and understand fundamental natural mechanisms, to replicate biological components, and ultimately, to engineer robotic systems that exhibit parallel functionalities. Recent strides in soft robotics have been pivotal in emulating the functionality of natural systems through the use of biomimetic materials (Coyle et al., 2018; Sachyani Keneth et al., 2021). These soft robotic systems offer distinct advantages, including ease of fabrication, inherent safety, and the ability to interact delicately with their environment or navigate through complex terrains (Shintake et al., 2018; Shepherd et al., 2011). Various modes of actuation such as elastomeric actuators, hydrogels, shape memory alloys (SMA), and electroactive polymers (EAP) have emerged as the mainstay in these systems (Appiah et al., 2019).

A focal point in soft robotics is the study of aquatic locomotion, a domain where nature exhibits remarkable agility and efficiency (Lauder et al., 2007). Fish, in particular, are renowned for their swift and efficient movement through dynamically changing aquatic environments (see Supporting Video 1), a trait that enables them to undertake strenuous tasks such as upstream migration during fasting periods (Crossin et al., 2004). The key to their exceptional energy efficiency lies in their ability to modulate the amplitude and frequency of their body undulations, leveraging the stiffness of their structure (McHenry et al., 1995; Lauder et al., 2011). By synchronizing body undulation frequency with the flow, fish swim efficiently, harnessing energy from the fluid and converting it into propulsion (Beal et al., 2006; Akanyeti et al., 2016; Liao and Akanyeti, 2017). The oscillation of the caudal fin emerges as one of the most efficient propulsion modes in terms of transport costs (Rayner, 1986; Ludeke and Iwasaki, 2019), and underwater speeds are enhanced significantly (Block et al., 1992), resulting in passive propulsion even in deceased fish specimens (Liao et al., 2003).

Research into fish locomotion have spawned a plethora of soft robotic designs capable of aquatic movement (Struebig et al., 2020; Nguyen and Ho, 2021). These designs range from robots that mimic the motion of tuna, surpassing their predatory speeds (Barrett, 1996; Zhu et al., 2019), to platforms that execute rapid ‘C-start’ maneuvers similar to carangiform fish (Marchese et al., 2014), and even robots capable of three-dimensional acoustic maneuvering (Hsieh et al., 2016; Katzschmann et al., 2018).Additionally, lateral body motions and reflex-based jumping capabilities have been integrated into robotic designs (Fan et al., 2005; Zhao et al., 2020; Kim et al., 2020; Yang et al., 2021). The intricacies of fish locomotion, particularly the interplay between active and passive stiffness control and the internal dynamics, offer a rich terrain for biomimetic technology transfer (Low and Chong, 2010; Low et al., 2010). Although fully passive fin systems have been developed to mimic fish propulsion, challenges arise in adjusting thrust output and drag in response to variations in flow velocity or frequency (Jayne and Lauder, 1996; Yun et al., 2011, 2015). To emulate the fish’s ability to adaptively modulate swimming body undulations and utilize soft surfaces, a sensory-driven control mechanism becomes essential.

Bio-inspired robotics opens up new possibilities for narrowing the gap between synthetic robots and their natural analogues (Hammond et al., 2023). This approach not only advances robotics but also provides biologists with tools to test hypotheses that may be impractical with living specimens, thereby serving as invaluable assets in biomechanical research (Siddall et al., 2021).

In our research, we draw inspiration from the locomotive prowess of rainbow trouts. Utilizing a soft robotic fish platform equipped with pneumatically actuated soft actuators and a soft bending sensor, as illustrated in Figure 1, we aim to advance the understanding and application of bio-inspired robotics. The soft bending sensor enables real-time monitoring and control of the bending amplitude of the robotic fishtail during oscillation. This setup is particularly suited for testing Repetitive Learning Controllers (RLC) (Xu and Yan, 2006; Marino et al., 2012; Verrelli, 2016, 2022). In our experimental approach, we implement a finite memory version of the RLC, using a method by (Tomei and Verrelli, 2015; Verrelli et al., 2015) to approximate delays in the learning estimation scheme with rational proper functions, leading to the development of the Linear Repetitive Learning Estimation Scheme (LRLES). This study presents a comparative analysis of trajectory tracking for a desired bending amplitude of the fishtail under constant oscillating frequency, using both a simple PID controller and the innovative PID-LRLES.

**Figure 1.**
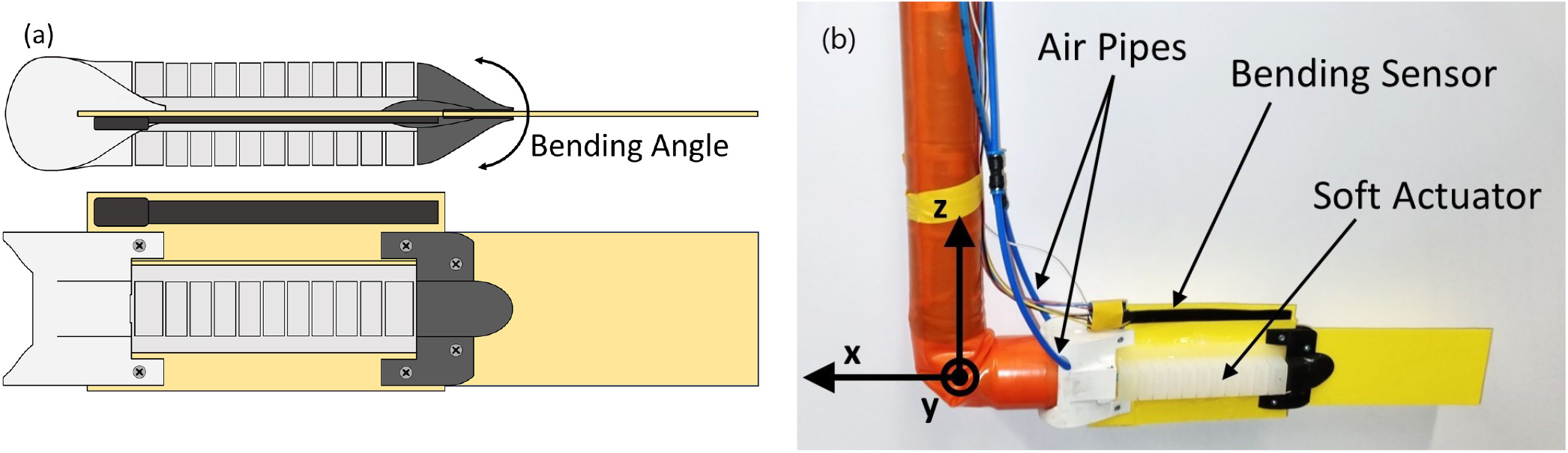
(a) Illustration of the soft pneumatic fish and (b) its physical realization.

The remainder of this manuscript is organized as follows: Section 2 outlines the experimental setup, the PID controller, and the PID-LRLES, along with the experimental procedures; Section 3 presents our experimental findings; and Section 4 discusses the implications of this work and outlines future research directions.

## 2 MATERIALS AND METHODS

### 2.1 Experimental Setup

#### 2.1.1 Soft Robotic Fish Platform

The soft robotic fish platform used in this work (Figure 1) consists of a flexible plastic foil, representing the backbone of the animal, to which two soft pneumatic actuators are glued to provide bending actuation. A frontal cuff (3D-printed, ABS) serves as an attachment point to the fixed mast, while another flexible tail cuff (3D-printed, TPU A95) holds the passive caudal fin. The cuffs also help streamline the overall fish profile. The main body is made following a similar method as described by Jusufi et al. (2017) and based on the work of Mosadegh et al. (2014). The flexible backbone foil (0.52 mm thick shim stock: Artus, Inc.) has a flexural stiffness comparable to that of a fish (9.9e-4 [Nm^2^]), and the actuators, made using a silicone-based elastomer (Dragon Skin™ 20, Smooth-On Inc.), are glued using a dedicated silicone adhesive (Dowsil 734 Flowable Sealant, DOW). A soft capacitive sensor (1-Axis Soft Flex Sensor, Nitto Bend Technologies) is affixed to a flexible foil extension above the midline to measure the bending angle while the fish is actuated. Care is taken to have both ends of the sensor rest above both cuffs to measure the bending angle across the whole body. Previous work from Wright et al. (2019) employed resistive eutectic gallium-indium (eGaIn) sensors to estimate the bending angle. As those sensors cannot measure both positive and negative angle values alone, two units, one on each side of the fish, have to be combined to extract a complete measurement. Additionally, for the bending to create sufficient stretching of the eGaIn sensors, they have to be placed at a distance from the midline. The capacitive sensor on the other hand can measure both positive and negative angles accurately, can be placed directly on the midline, and requires less calibration effort. For these reasons, and because the resistive sensors are also more fragile, the capacitive sensor was selected in the end.

The pressure network system used to power the soft actuators is illustrated in Figure 2. A digital pressure regulator (ITV0050-3BS, SMC) supplies between 0.7 and 2.5 bar (actuator working range) of compressed air to the system in a controlled fashion. Each actuator is governed by two solenoid valves (SYJ7320-5LOU-01F-Q, SMC) that either connect it to the pressure system controlled by the regulator, or the atmosphere. An air tank allows to store some pressure during actuation and helps mitigate the variations created by the release of actuator pressure in the atmosphere. A real-time microcontroller (myRIO-1900, National Instruments) acts as the Control Unit of the robotic platform. It samples the bending sensor via I2C at 1kHz and runs the control loop at the same frequency, generating the actuating signals to the valves and control signal to the pressure regulator as displayed in Figure 3.

**Figure 2.**
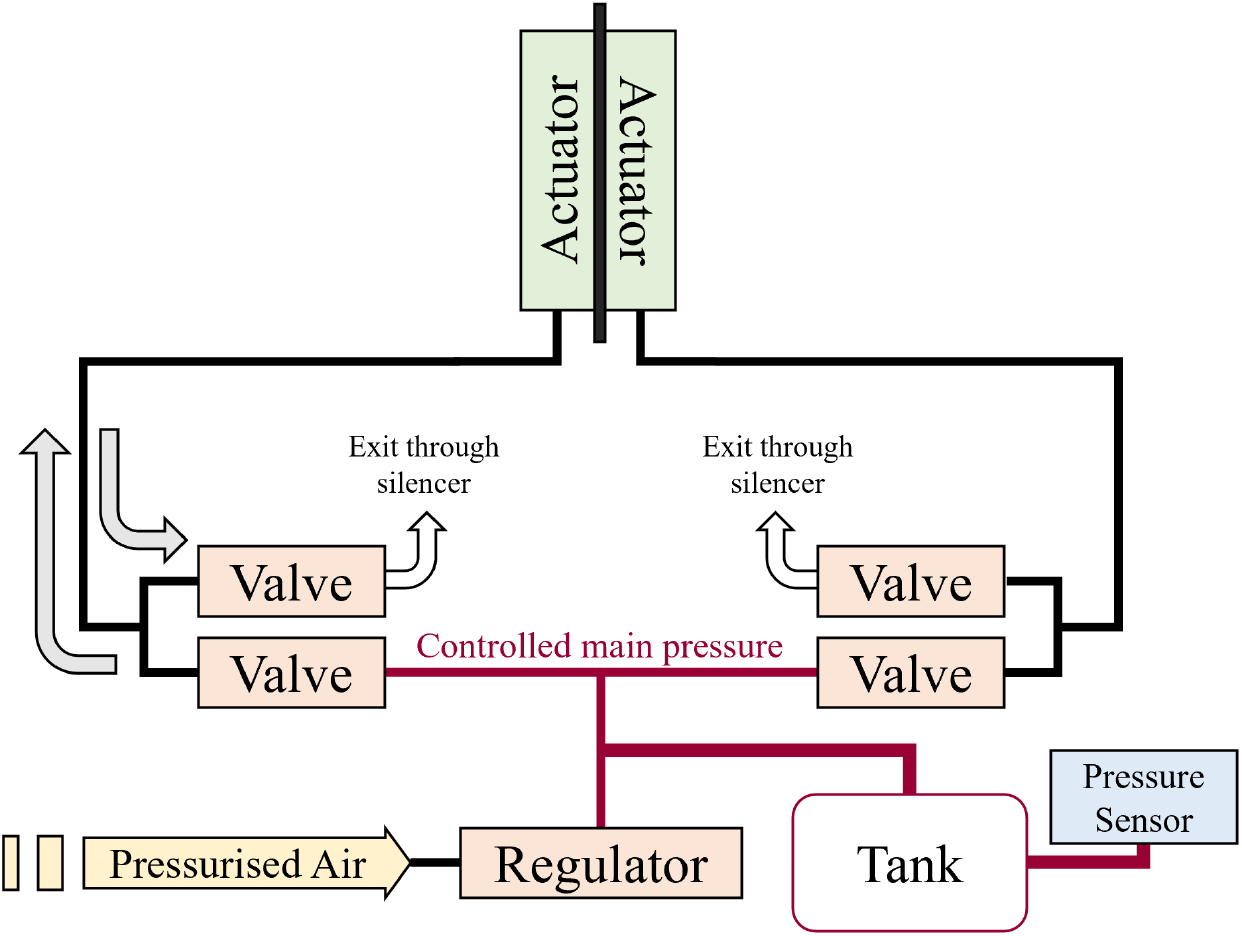
Schematic of the soft fish platform’s pneumatic actuation circuit. The external pressurized air source provides the regulator with a stable 2.5 bar input. For each actuator at any point in time, one of the two connected solenoid valves is opened to either fill the actuator at the controlled main pressure (0.7 to 2.5 bar) or release it to atmospheric pressure.

**Figure 3.**
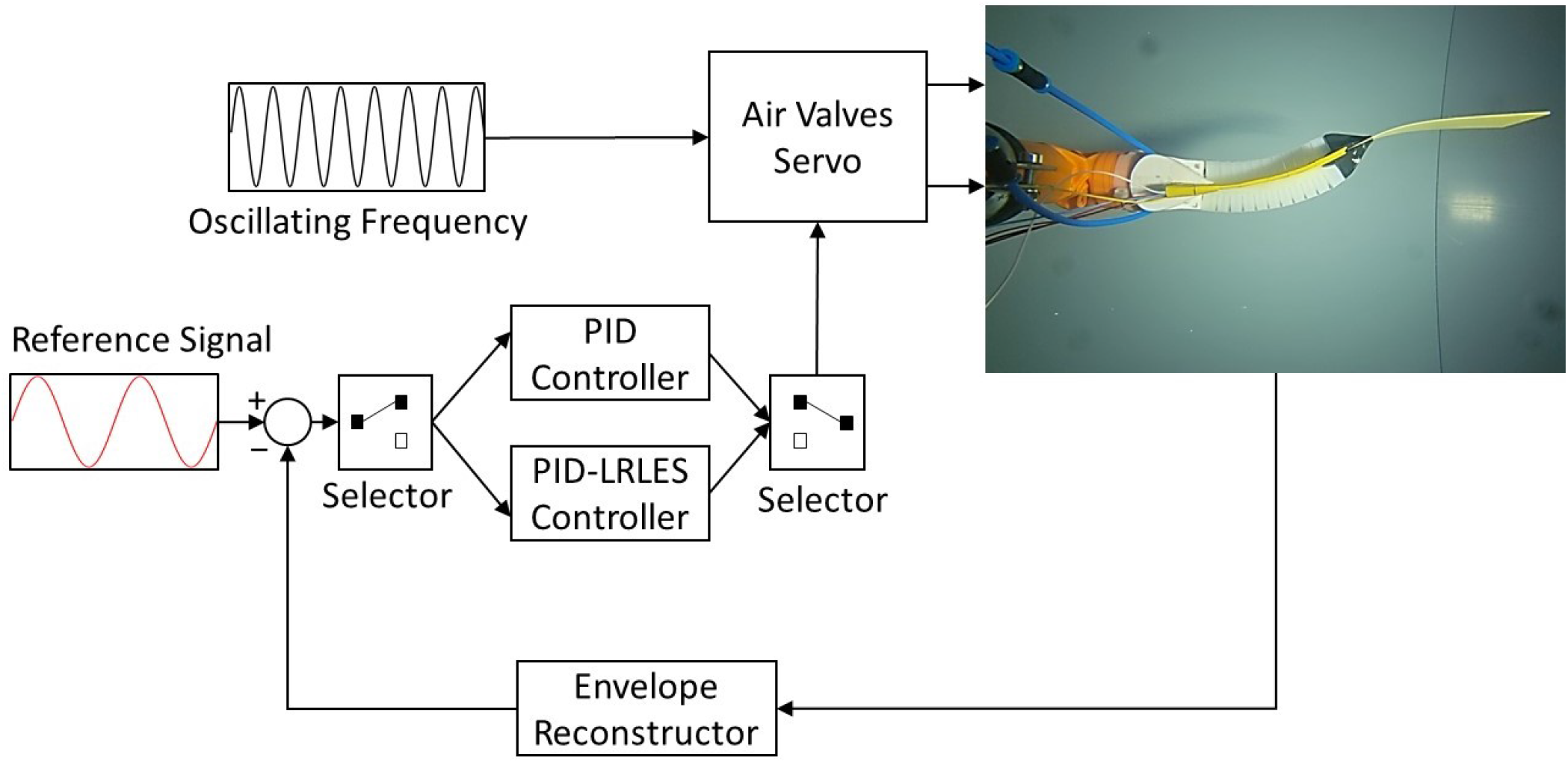
Illustration of the controller implemented on the MyRIO to control the soft robotic fish.

#### 2.1.2 Test Rig

Experiments were conducted within a large water tank located at the Swiss Federal Laboratories for Material Science (EmpaMPA) with a cross-sectional area measuring 0.6 *×* 1 m^2^, accommodating a fully transparent test section extending up to 6 m in length (Figure 4a).

**Figure 4.**
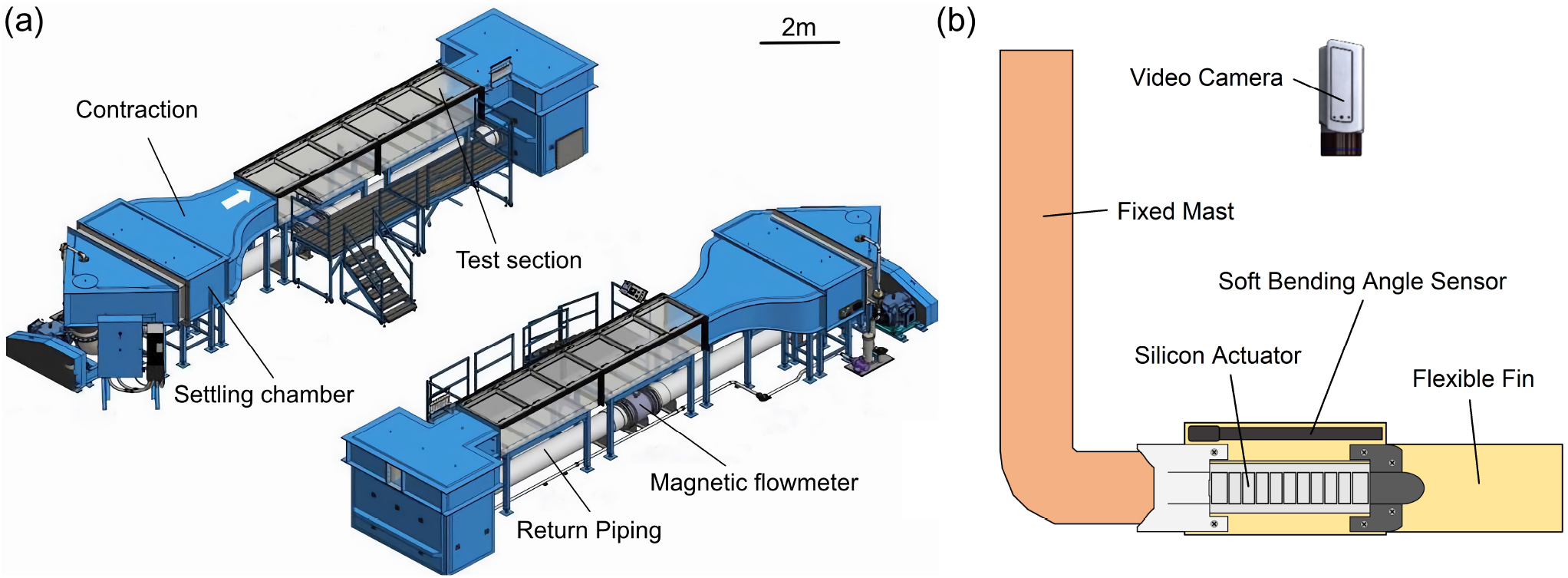
Experimental setup for the soft robotic swimming experiments. (a) General setup of the water tank. (b) Schematic of the robotic platform placed in the water flow tank.

The soft robotic fish platform was suspended in the center of the test section and secured in place by two extra aluminum beams on both sides of the mast for stability. A video camera was placed above the platform to capture the robotic fish’s midline kinematics (Figure 4b). To avoid waves and reflections at the water surface, the recording was made through a Plexiglas tray in contact with the water at all times.

### 2.2 From PID control to PID-LRLES

The control of the soft robotic fish aims at directing and effectively deploying an untethered underwater robot mimicking the size and behavioral traits of living fish. Controlling robotic fish plays a pivotal role in achieving biomimetic swimming performance, ultimately leading to a system that closely resembles its natural counterparts. While various model-based control approaches can potentially be applied to soft robotic systems akin to other robotic platforms, the lumped nature of soft-system models – with internalstates just being representative of complex neglected dynamics and parameters that are unfortunately uncertain – limits the applicability of certain theories and algorithms that require more than the knowledge of the structural properties of the model, such as relative degree, high-frequency gain, and minimum-phase. Controlling soft robots thus presents a significant challenge, necessitating the exploration of viable and optimally effective control strategies capable of managing the soft body and leveraging its intrinsic intelligence.

In our model, the primary control objective revolves around regulating the pressure within the soft actuators. This pressure modulation dynamically influences the entire robotic fish system by altering thrust and side force, generating torque at each joint, ultimately impacting the final bending angle of the tail, considered as an end-effector of the robot.

Owing to the repetitive features of the body caudal fin undulation of the soft robotic fish, the most suitable control design is one that solves an output tracking control problem for uncertain nonlinear systems characterized by unstructured uncertainties (that is when no parameterization of the uncertainties is available) and by output reference signals that belong to the family of periodic time functions with a known period (piece-wise constant references are just special cases). RLCs are good candidates (Marino et al., 2012; Verrelli, 2016, 2022), since they are not model-based and aim at performing a dynamic system inversion just through the feedback. However, finite memory implementations of control laws are needed in practice. They should involve the use of a finite number of stored values or be finite-dynamic-order controls. This is the case of the linear repetitive learning control of (Verrelli et al., 2015) (see theoretical foundations in (Tomei and Verrelli, 2015)), in which the delay involved within the repetitive learning estimation scheme is approximated by Padé theory-based rational proper functions, so as to approximate a delay-based infinite-dimensional system by a finite-dimensional one. Owing to the dynamic linearity of Padé approximants and to the use of a stabilizing filter, the LRLES generalizes the classical integral actions (typically used within the PID control for robotic systems, with no “zeros” and relative degree two) to the case of periodic (non-constant) references: they are described by transfer functions the poles of which all have a negative real part such that the typical long-term instability issues of classical RLCs, due to high-frequency disturbance noises, are avoided, thus leading to strong stability properties.

In this section, a LRLES of such a type (referred to as ℒ 𝒞 (*s*) hereafter) is added to the control scheme for fish robots, to endow the PID control with a mechanism that is able to track periodic reference signals besides constant ones. Referring to the model established in (Lin et al., 2021), where the design was applied to an *n*-link rigid structure, *M*_*n*_ represents the vector consisting of the control torques of the pressure inside the fish actuator. The control problem thus concerns the design of the controller for the last joint *q*, which will provide the maximum bending angle of the fish due to the end of the tail (referring to Figure 5 (a), *q* is obtained by taking the modulus of the bending angle and computing its envelope). The scenario is thus similar to the one in (Verrelli et al., 2015), once the feedback signal,

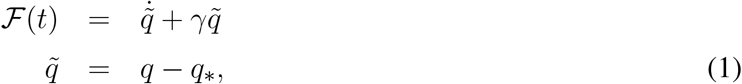

is defined by the (possibly filtered) linear combination 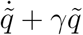 (with positive weights *γ* and 1) of the joint angle tracking error 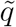 and its time derivative 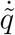, where *q* denotes the available output and *q*_*_ the periodic output reference signal.

**Figure 5.**
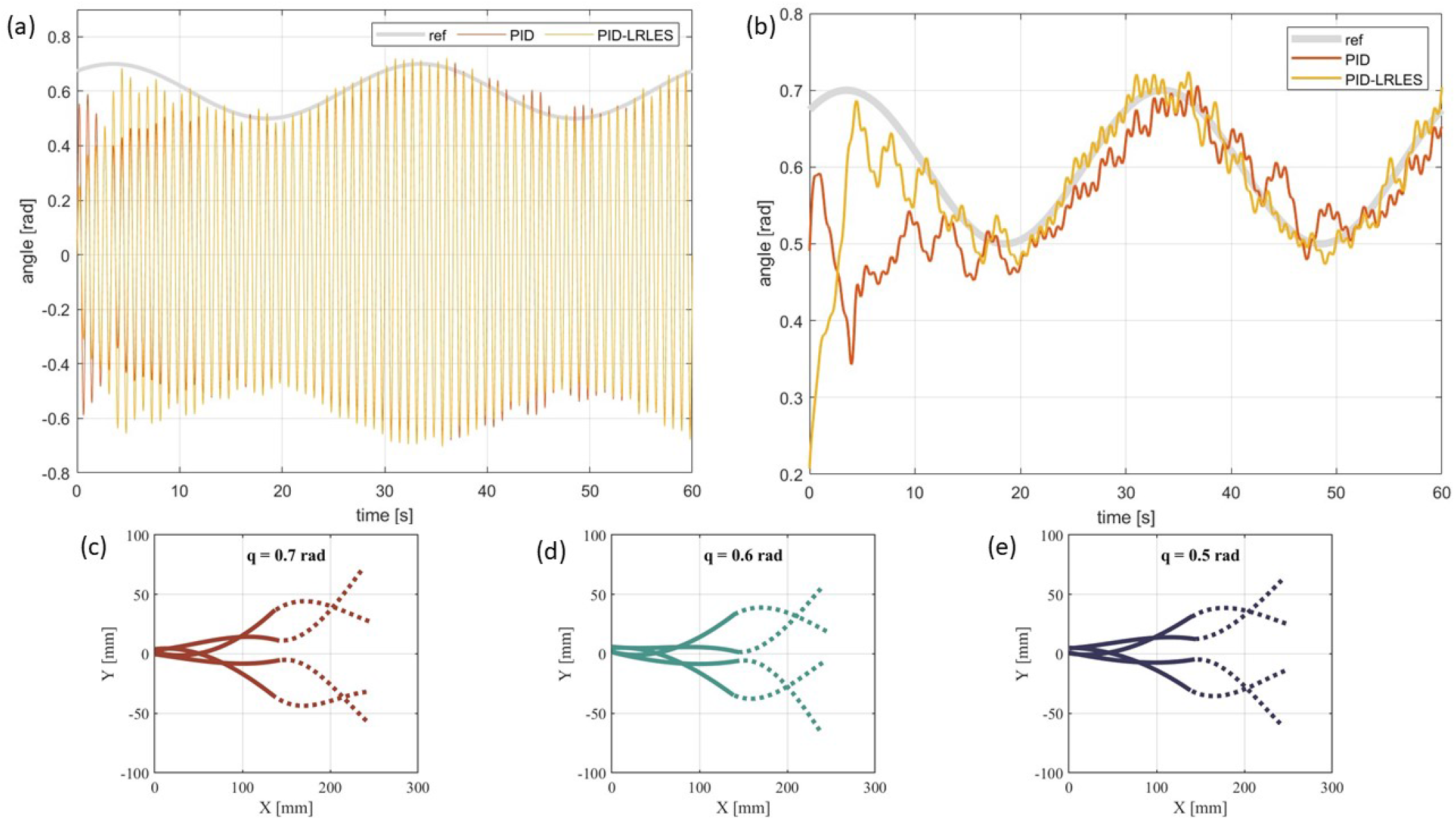
The extract analyzes the tracking accuracy of a soft robotic fish platform, specifically comparing the bending angle (a) and the envelope (b) under different control conditions. ‘ref’ denotes the target amplitude envelope for the tail oscillation. ‘PID’ represents the data measured using the Proportional-Integral-Derivative (PID) controller. ‘PID-LRLES’ refers to measurements obtained using a PID controller augmented with the Linear Repetitive Learning Estimation Scheme (LRLES) method. (c) to (e) illustrate the midline motion pattern of the fish’s foil during a complete swimming swing, displaying variations in oscillation magnitude while tracking signals with different bending amplitude envelopes *q*. The thick lines indicate the backbone of the fish, where the bending angle is measured, while the dashed lines represent the passive movement of the tail. The leading edges are on the left.

#### 2.2.1 PID control

The PID controller,

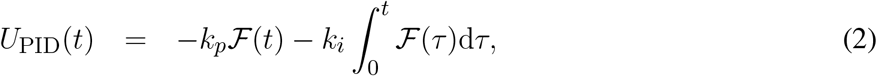

with transfer function

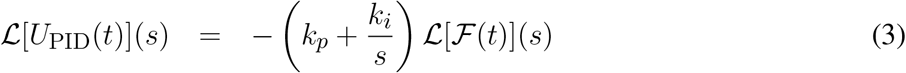

was initially employed for the control of a soft robotic fish by (Lin et al., 2021). If 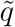 is the only available error (and the derivative can be numerically obtained), then the previous expression can be rewritten for zero initial conditions as

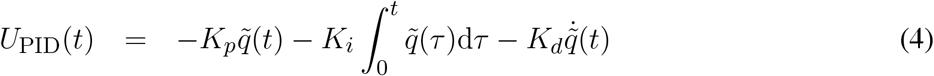

in terms of the proportional, integral, and derivative gains

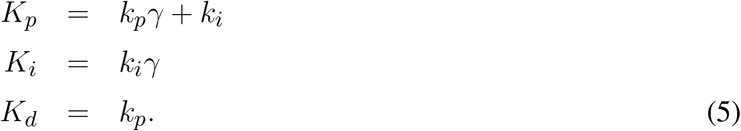

This controller is widely used in diverse dynamic systems, including industrial, robotic, and biological applications and stands as the first viable candidate due to its robust performance. The PID controller operates by continuously calculating the error value from the difference between a desired setpoint and the measured process variable. It incorporates a proportional and a derivative term – each with the corresponding gain – to stabilize the error system, along with an integral term to reconstruct the constant reference input that guarantees perfect output regulation to any constant output reference.

An amplitude control system, prototyped in Simulink, incorporating sensor noise effects, was designed and tested (Lin et al., 2021). This control system extracts the maximum amplitude within each half period from the oscillating strain signal. The PID controller demonstrates robust behavior against sensor noise, exhibiting fluctuations around the step point under noises. Furthermore, periodic references, though piece-wise constant, repeatedly induce transient error behaviors.

#### 2.2.2 PID-LRLES

The RLC idea comes from the observation that animals can enhance task execution through repeated trials. In contrast to non-learning controllers, the repetitive learning mechanism utilizes the error signal coming from *p* previous executions (*p ∈* ℕ^+^) within a repetitive framework in order to harness experience and refine the closed-loop performance.

Following (Tomei and Verrelli, 2015), let the [*m, m*]-Padé approximant of e^*−sT*^ being given by

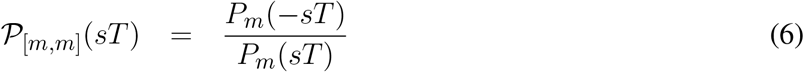

with

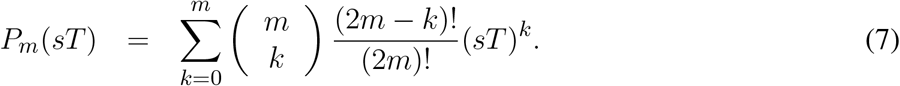

Then define the proper transfer function

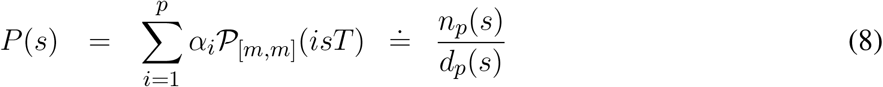

which approximates the Laplace transform 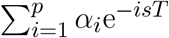 of the delay mechanism contained within the “high-order” repetitive learning estimation scheme of (Tomei and Verrelli, 2015):

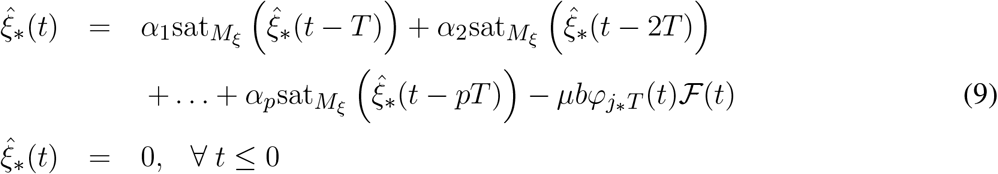

in which:

- 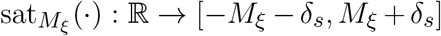 is a class 𝒞^1^ odd increasing function satisfying (*δ*_*s*_ is an arbitrary positive real) 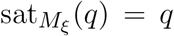 for any 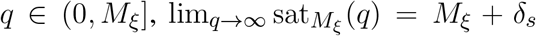 and 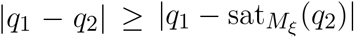 for any |*q*_1_| *≤ M*_*ξ*_, *q*_2_ *∈* ℝ;
- 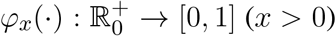 is a class 𝒞^1^ increasing function for *t ∈* [0, *x*] with 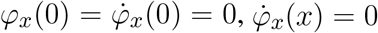 and *φ*_*x*_(*t*) = 1 for any *t ≥ x*, which endows the 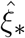-signal with some continuity properties;
- *μ* and *α*_*i*_, 1 *≤ i ≤ p*, are positive design parameters with *α*_*i*_ satisfying

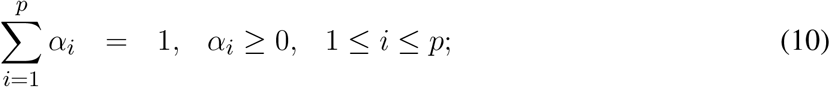

*b* = *α*_1_ + 2*α*_2_ + … + *pα*_*p*_ and *j*_*\**_ = min*{j* : *α*_*j*_ *>* 0*}*.

Such a learning estimation scheme relies on a weighted sum of the information stored in the *p* previous executions: the weights *α*_*i*_, 1 *≤ i ≤ p* are extra degrees of freedom that allow the control design to take into account the whole available information about the past to improve the performance of a periodic system. If 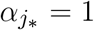 and *α*_*i*_ = 0 for 1 *≤ i ≤ p* with *i* ≠ *j*_*\**_, then the estimation scheme reduces to the classical ‘first order’ one in which the input reference *ξ*_*\**_(*t*) is interpreted as a periodic signal with period *j*_*\**_*T*.

When the action of the saturation function is neglected and *φ*_*j\**_*T* (*t*) *≡* 1 is considered, (9) reduces to

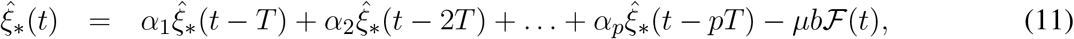

which satisfies

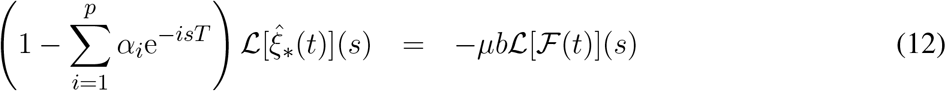

in the Laplace domain. On the other hand, if *ξ*_*\**0_(*t*) denotes the function which is equal to *ξ*_*\**_(*t*) on the set [0, *T*) while being zero outside it, then we can write

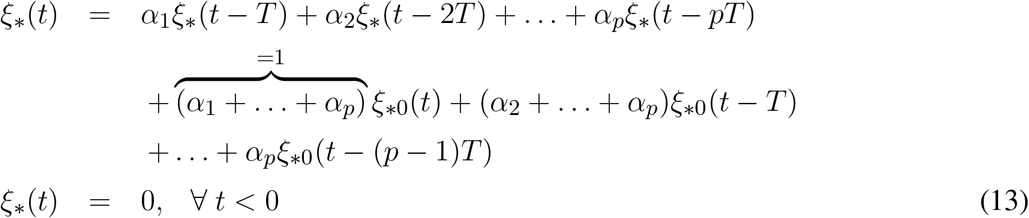

which plays the role of an infinite-dimensional exosystem reproducing the time *T* -periodicity of *ξ*_*\**_(*t*). The initial function *ξ*_*\**0_(*t*) (just like the initial condition for a finite-dimensional exosystem) is the only unknown quantity to the controller to be dealt with by means of the feedback action *−μbφ*_*j\**_*T* (*t*) ℱ (*t*). We then rewrite the above exosystem as

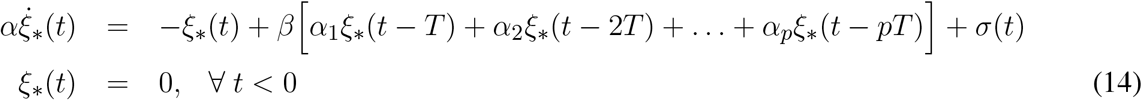

where the signal *σ*(*t*) can be obtained by comparison. Then, we design the (1 + *p · m*)-finite-dimensional approximation of the exosystem as 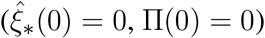

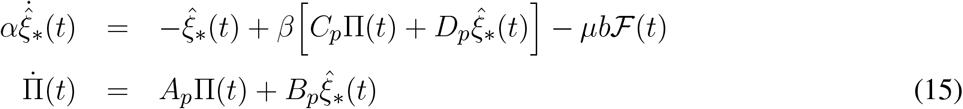

where (*A*_*p*_, *B*_*p*_, *C*_*p*_, *D*_*p*_) is a minimal realization of the proper transfer function *P* (*s*). Here, *A*_*p*_ is a Hurwitz matrix, *d*_*p*_(*s*) is a *p · m*-order polynomial whereas *α ∈* [0, 1), *β ∈* (0, 1] are assumed to guarantee that the polynomial

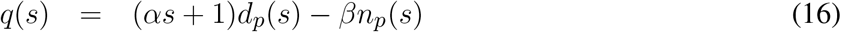

has all the roots belonging to ℂ^*−*^. The term 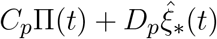 is nothing else than the Padé approximation of the delayed term 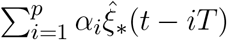. The resulting filter *β/*(*αs* + 1) is introduced to force the learning estimation scheme in the Laplace domain

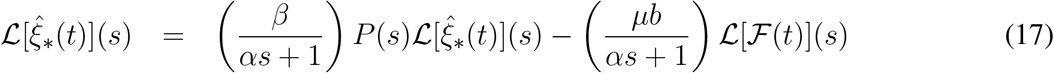

to have a transfer function

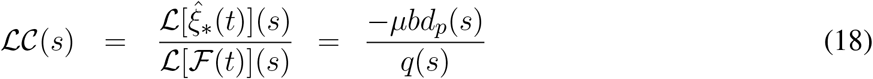

with all the poles belonging to ℂ^*−*^. Such a transfer function reinforces the integral action within the controller, leading to the PID-LRLES:

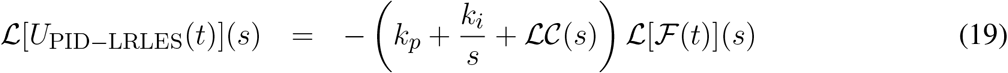

The incorporation of the (*α, β*)-filter is necessary for this purpose since, when *α* = 0 and *β* = 1, *s* = 0 is a root of *q*(*s*) according to

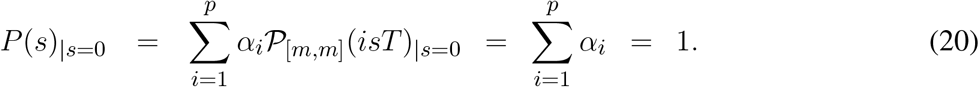

Notice that, for *α* = 0, *β <* 1 and *α*_1_ = 1, *α*_*j*_ = 0 (*j* = 2, …, *p*), the equivalent exosystem

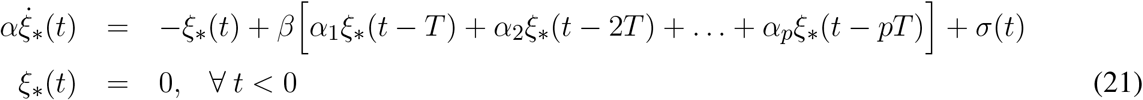

becomes

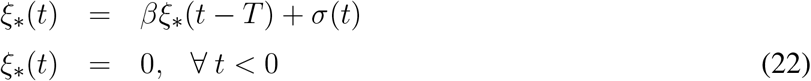

where the solution of

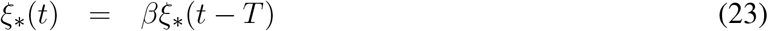

evaluated at any *t* = *t*_*\**_ + *kT* (*t*_*\**_ *∈* [0, *T*), *k* = 0, 1, …) constitutes a decreasing sequence converging to zero. The preceding equation can be rewritten as

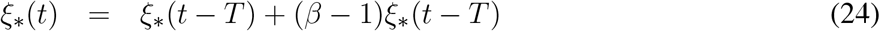

in which the second term on the right-hand-side (with *β −* 1 sufficiently small) plays a stabilizing role, which leads to a bounded-input bounded-output estimation law without the use of saturation functions or projection algorithms.

### 2.3 Experimental Procedure

The experiment consists of tracking a desired trajectory for the envelope containing the amplitude of the bending angle of the robotic fish. The platform is actuated at a constant frequency of 1.2 Hz and at 0.55 duty cycle (5% co-contraction). The envelope is used to obtain a positive, smoother signal. The desired amplitude of the envelope is described by a sine wave with period *T* = 30 s, an amplitude of 0.1 rad, and a 0.6 offset. The experimental procedure compares the tracking performance of the PID controller and the PID-LRLES. Both controllers have been separately fine-tuned manually during the experiments, with the PID controller gains *K*_*p*_ = 3, *K*_*i*_ = 0.28, and *K*_*d*_ = 0.15 and the PID-LRLES gains *m* = 7, *p* = 3, *α*_1_ = 0.3, *α*_2_ = 0.2, *α*_3_ = 0.5, *γ* = 4, *β* = 0.99, *μ* = 0.4950, *k*_*i*_ = 0.3 and *k*_*p*_ = 0.01. Notice that the PID controller, which does not contain the learning estimation scheme, has to increase its proportional and derivative gains – compared to the ones within the PID-LRLES (recall (5)) – and decrease the integral gain in order to exhibit a comparable – though worse – closed-loop behaviour.

All experiments start with the fish oscillating in an open loop, meaning that the actuation chambers are inflated alternatively for a fixed and predefined amount of time. After some iterations, the amplitude of the bending angle envelope reaches a steady state, and at that moment, the desired controller is turned on, allowing the robotic fish to track the desired trajectory for the amplitude of the bending angle envelope.

## 3 RESULTS

The experimental procedure compares the performance of the PID controller and the PID-LRLES in tracking a desired trajectory of the fishtail bending angle envelope. Figure 5.(a-b) shows the tracking accuracy during the experiment, which lasts for 60 seconds (two periods of the sinusoidal reference signal). The PID-LRLES performs better than the PID controller in tracking the desired trajectory (though the PID controller was separately tuned), being at the same time faster and more accurate. The root mean square error (RMSE) is calculated over an oscillation period of the reference trajectory (20-50 s). The PID-LRLES outperforms the PID controller, the RMSEs being respectively *E*_PID*−*LRLES_ = 0.015 rad and *E*_PID_ = 0.032 rad. The right side of Figure 5 (d-e) illustrates the fish kinematics by plotting the mid-line kinematics of the fish backbone (solid line) and tail (dashed line) in three different moments of the experiments, with the fish tracking respectively the minimum, the medium, and the maximum tracked values for the bending angle envelopes *q* = 0.5, *q* = 0.6 and *q* = 0.7.

The high-frequency oscillations in Figure 5.b are at 1.2 Hz, which is the oscillating frequency of the fish. We attribute this behavior to the abrupt changes in the oscillatory direction that happen when the control system switches the active actuator, causing the 3D-printed support to rattle in its location (see accompanying video). Both controllers proved to be robust to unmodeled disturbance at a price of small oscillations.

## 4 DISCUSSION

In this study, we have successfully developed and implemented a Proportional-Integral-Derivative with a Linear Repetitive Learning Estimation Scheme (PID-LRLES) controller on a soft robotic fish platform. This novel controller demonstrates a significant improvement in performance over the traditional PID controller in tracking periodic signals. Its success is particularly notable in the context of soft robotic systems, which often contend with uncertain parameters and unmodeled dynamics. This advancement is not just a technical achievement but also a critical step forward in the field of soft robotics, showcasing the potential of advanced control methods in managing complex dynamic systems.

Our work builds upon and extends our previous foundational studies (Jusufi et al. (2017), Lin et al. (2021) and Schwab et al. (2022)). While Jusufi et al. focused on open-loop control experiments without feedback on the oscillating tail’s position, and Lin et al. limited their exploration to a PID controller for single step-response tracking, our approach achieves faster and more accurate tracking. This progress is attributed to both the innovative PID-LRLES controller and enhancements in the mechanical structure of the robotic fish. However, the pioneering work by Lin et al. remains a valuable reference, especially in their use of eutectic gallium-indium (eGaIn) stretch sensors for bending angle measurements.

While our controller’s performance in environments with incoming water flow remains untested due to unforeseen constraints, we hypothesize that the PID-LRLES controller could effectively estimate and compensate for drag variations caused by flow. This capability would mark a significant advancement in robustness compared to classical control methods. Future research will focus on employing resistive eGaIn soft sensors to test this hypothesis. These sensors are expected to mitigate the electromagnetic disturbances encountered in the flow tank and refine the process of bending angle measurement.

Looking ahead, our aim is to further develop the soft robotic fish platform, with a specific focus on experimental validation in flow tank conditions. The next phase of research will involve integrating the flow regime into the control structure of the PID-LRLES, aiming for a more holistic and responsive control system. This approach promises to open new avenues in the design and control of soft robotic systems, contributing to our understanding of biomimetic robotics and its application in dynamic aquatic environments.

## ACKNOWLEDGMENTS

We thank the Central Service Station for Robotics at the Max Planck Institute for Intelligent Systems for practical support. We thank Bingcheng Wang of Soft Kinetic Group for the kinematics analysis of undulatory body caudal fin swimming. We thank Roger Vonbank of the Laboratory for Multiscale Studies in Building Physics for his support with the experimental setup.

## SUPPLEMENTAL DATA

Supplementary Material includes a video file accompanying the paper where the first video sequence is copyrighted by Journal of Fluid Mechanics from Beal et al. 2006 and the second video is copyrighted by Soft Robotics Journal from Jusufi et al. 2017.

## DATA AVAILABILITY STATEMENT

The datasets for this study can be found here https://github.com/ardianet1/SoftSensoRepetLearn. Other data presented in this paper can be made available upon reasonable request.

